# PIC recruitment by synthetic reader-actuators to polycomb-silenced genes blocks triple-negative breast cancer invasion

**DOI:** 10.1101/2023.01.23.525196

**Authors:** Natecia L. Williams, Lauren Hong, Maya Jaffe, Cara E. Shields, Karmella A. Haynes

## Abstract

Scientists have used small molecule inhibitors and genetic knockdown of gene-silencing polycomb repressive complexes (PRC1/2) to determine if restoring the expression of tumor suppressor genes can block proliferation and invasion of cancer cells. A major limitation of this approach is that inhibitors can not restore key transcriptional activators that are mutated in many cancers, such as p53 and members of the BRAF SWI/SNF complex. Furthermore, small molecule inhibitors can alter the activity of, rather than inhibit, the polycomb enzyme EZH2. While chromatin has been shown to play a major role in gene regulation in cancer, poor clinical results for polycomb chromatin-targeting therapies for diseases like triple-negative breast cancer (TNBC) could discourage further development of this emerging avenue for treatment. To overcome the limitations of inhibiting polycomb to study epigenetic regulation, we developed an engineered chromatin protein to manipulate transcription. The synthetic reader-actuator (SRA) is a fusion protein that directly binds a target chromatin modification and regulates gene expression. Here, we report the activity of an SRA built from polycomb chromodomain and VP64 modules that bind H3K27me3 and subunits of the Mediator complex, respectively. In SRA-expressing BT-549 cells, we identified 122 upregulated differentially expressed genes (UpDEGs, ≥ 2-fold activation, adjusted *p* < 0.05). On-target epigenetic regulation was determined by identifying UpDEGs at H3K27me3-enriched, closed chromatin. SRA activity induced activation of genes involved in cell death, cell cycle arrest, and the inhibition of migration and invasion. SRA-expressing BT-549 cells showed reduced spheroid size in Matrigel over time, loss of invasion, and activation of apoptosis. These results show that Mediator-recruiting regulators broadly targeted to silenced chromatin activate silenced tumor suppressor genes and stimulate anti-cancer phenotypes. Therefore further development of gene-activating epigenetic therapies might benefit TNBC patients.

## INTRODUCTION

The concept of epigenetic cancer therapy is based on the important discovery that many anticancer genes are not mutated, but instead lie dormant in an epigenetically repressed state. Determining whether and how transcription can be restored at these domain genes is potentially transformative for cancer therapy. At transcriptionally activated genes, the Mediator complex coordinates the assembly of the preinitiation complex (PIC) at the promoter. One major mechanism for gene silencing in cancer is the assembly of polycomb complexes at enhancer and promoter regions to form a transcriptional blockade (**Fig. 1A**). Polycomb repressive complex 2 (PRC2) contains EZH2, which methylates histone H3 lysine 27. H3K27me3 acts as a docking site for the CBX subunit (CBX2, 4, 6, 7 or 8) of canonical polycomb repressive complex 1 (cPRC1) which ubiquitylates histone H2AK118 and K119^1^. An *in vitro* study showed that PRC1 disassembles PIC by evicting Mediator proteins while TBP, a member of the TFIID complex, remains bound^2^. In mouse embryonic stem cells, genes that are occupied by TBP and PRC1 are expressed at lower levels than genes where Mediator is also present.

**Figure 1.**
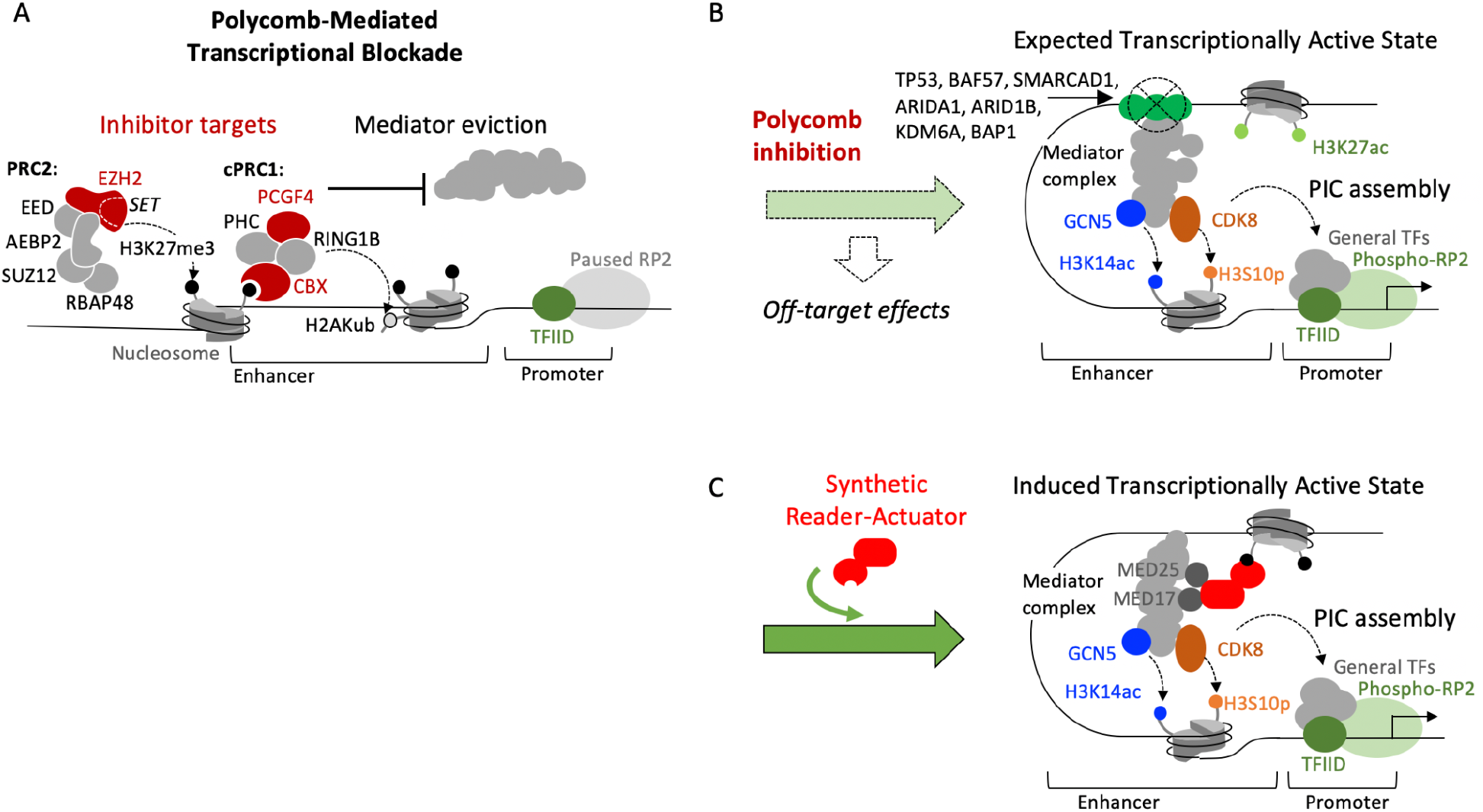
A synthetic reader-actuator designed to stimulate PIC assembly at polycomb-repressed genes. (A) Model for epigenetic repression of genes by hyper-active polycomb proteins. Targets of small molecule inhibitors are highlighted. (B) Transcription factors that may be important for gene reactivation (green circles), but are often mutated in cancer, are listed. (C) Model for SRA-mediated transcriptional activation. TF, transcription factor; RP2, RNA polymerase II; Phospho-RP2, phosphorylated RP2.

Elevated levels of polycomb proteins EZH2, BMI1/*PCGF4*, RING1B/*RNF2*, and CBX2 in basal-like and triple-negative breast cancer (TNBC) tumors may be responsible for silencing genes that block cancer cell proliferation and invasion^3–5^. Overexpression of EZH2 was determined to have strong prognostic value, i.e. prediction of metastasis within five years of primary diagnosis and death, above and beyond other biomarkers in 2003^3^. In studies of tumors from patient cohorts (42 to 280 individuals), abnormally high levels of EZH2, BMI1, and CBX2 correlate with more aggressive TNBC, i.e. higher tumor grade, recurrence after initial treatment, metastasis, and low survival^3–8^. Epigenetic activation of tumor suppressor genes represents a potential new therapeutic approach for hard to treat solid cancers such as TNBC. Nearly 80% of TNBC patients who receive neoadjuvant chemotherapy experience recurrence and metastases to the brain and lungs, which occur more often and are deadlier than for other types of breast cancer^9,10^. The prevalence of TNBC in African-American women^11–14^ facing healthcare disparities further compounds the devastating impact of this disease. The alarming failure rate for therapy against TNBC has been linked to cellular resistance against cell cycle arrest and apoptosis^6,15,16^.

Removal of polycomb repressive complexes (PRCs) from DNA by pharmacological inhibition or genetic knockdown is expected to allow assembly of the preinitiation complex (PIC) at promoters. However, with this approach there is no step to recruit and install transcriptional activators. Studies using mouse embryonic stem cells showed that depletion of either PRC1 or PRC2 did not increase DNA accessibility at polycomb-occupied genes^17,18^ which suggests incomplete conversion to a transcriptionally active state. FDA-approved tazemetostat partially reduces the viability of triple-negative breast cancer cells *in vitro^19^* suggesting that polycomb inhibition does not fully activate apoptotic gene expression in TNBC. Dysfunctional transcription factors may pose a barrier to tumor suppressor gene activation after repressive chromatin proteins are removed (**Fig. 1B**). TP53 binds the ACID domain of MED25^20^ to support the transcription of many tumor suppressor genes, but TP53 is mutated (loss of DNA binding) in 80%-95% of TNBC cases^19,21^ and in cell lines BT-549 (R249S), MDA-MB-231 (R280K), and MDA-MB-468 (R273H)^22^. The human BAF SWI/SNF (switch/sucrose nonfermentable) chromatin remodeling complex can evict PRC1 from the tumor suppressor locus INK4b-ARF-INK4a, as shown in malignant rhabdoid tumor cells^23^. However a key SWI/SNF subunit BAF57 is not expressed in cell line BT-549^24^, and subunits SMARCAD1, ARID1A, ARID1B, KDM6A, and BAP1 are frequently mutated (35%) in mesenchymal tumors^19^. Furthermore, polycomb protein inhibition may induce off-target effects by disrupting non-PRC regulatory complexes that contain polycomb proteins (**Fig. 1B**).

We designed an engineered chromatin protein as a new way to probe transcriptional plasticity within polycomb-repressed chromatin. The SRA “Polycomb transcription factor” (SRA-PcTF) contains an N-terminal H3K27me3-binding polycomb chromodomain (PCD) from human CBX8 and a transcription initiation factor-binding domain (4xVP16, “VP64”) and a nuclear localization signal (NLS) (**Fig. 1C**). We have shown that the targeting module, PCD, specifically binds H3K27me3 (not K4me3 or K9me3) peptides *in vitro^25^*. The activation module contains four copies of VP16, and each has strong affinity for the ACID domain of MED25 and MED17 of the Mediator master regulator complex^26,27^. We showed that SRA-PcTF stimulated transcription at hundreds of genes, including known polycomb-repressed loci (e.g. *HOXB4*), and genes involved in cell cycle arrest (e.g. CDKN1A) and cell differentiation^28^ in osteosarcoma cells, and immunomodulatory genes in breast cancer cell lines^29^.

In this study, we identified 122 protein-coding genes that become activated by SRAs in triple negative breast cancer cells (BT-549) with elevated polycomb protein levels. When we tried to identify activatable H3K27me3-enriched genes in BT-549 cells by inhibiting specific polycomb complex subunits, we observed treatment-specific transcriptional changes in either direction (up or down-regulated) at lowly-expressed, highly-expressed, and H3K27me3-enriched and depleted genes. In contrast, the vast majority of differentially regulated genes in SRA-PcTF expressing cells were activated, and these target genes were enriched for H3K27me3. While activated genes belonged to both tumor suppressing and cancer promoting pathways, SRA-mediated transcriptional activation showed an anti-cancer phenotypic effect as demonstrated by reduced size of BT-549 spheroids in Matrigel over time, loss of invasion, and activation of apoptosis.

## RESULTS

### Standard polycomb-targeting approaches show inconsistent effects on transcription levels

Polycomb inhibition is expected to remove the polycomb blockade at promoters to allow transcription. We subjected BT-549 cells to genetic knock-down or pharmacological inhibition of core subunits of canonical cPRC1 and PRC2, and measured changes in gene expression with a Nanostring assay. EZH2, the catalytic subunit of PRC2, has a SET domain that methylates histone H3 lysine 27 and can be blocked by SAM-competitive compounds such as GSK126^30^ and GSK343^31^. PCGF4 (BMI1) is a core component of cPRC1 that supports the H2A ubiquitin ligase activity of RING1A/B (*RING1*)^32,33^, and engages in protein-protein interactions that stabilize the entire complex ^34^. Compounds PTC209 and PTC596 block translation of PCGF4^35^ and possibly trigger its degradation^36^. The CBX (chromobox) subunit of cPRC1 contains an evolutionarily-conserved N-terminal hydrophobic pocket that binds H3K27me3, and a C-terminal disordered region that supports assembly of the other cPRC1 subunits. The compound UNC3866 occupies the hydrophobic pocket of CBX4 and CBX7 to block binding with H3K27me3^37^.

To verify high polycomb protein levels, which support hyper-repression of target genes, we used public data (cBioPortal, CCLE Broad study) to compare the expression of genes that encode subunits of canonical polycomb complexes in TNBC versus non-cancer mammary epithelial HMEL cells. BT-549, MDA-MB-231, and MDA-MB-436 cell lines of the mesenchymal sub-class, which is invasive and drug-resistant^38^, showed the highest relative polycomb levels for PCGF4 and EZH2 (**Fig. 2A**). Using a western blot assay, we detected higher levels of EZH2 and PCGF4 in BT-549 compared with the non-cancer mammary epithelial cell line hTERT-HME1 (**Fig. 2B**). In a recent comparative study, mesenchymal TNBC cells showed the highest H3K27me3 levels, which is consistent with high EZH2 levels^19^. Our analysis of public RNA-seq data showed that mRNA levels for EZH2 and CBX2 were elevated close to 1.5-fold for paired normal/ basal TNBC patient samples (Xena database), and that tumor suppressor genes^39^ were downregulated in basal TNBC compared to normal breast samples (cBioPortal) (**Supplemental Fig. S1**), which supports the clinical significance of abnormally high levels of polycomb in aggressive breast cancer^3–8^.

**Figure 2.**
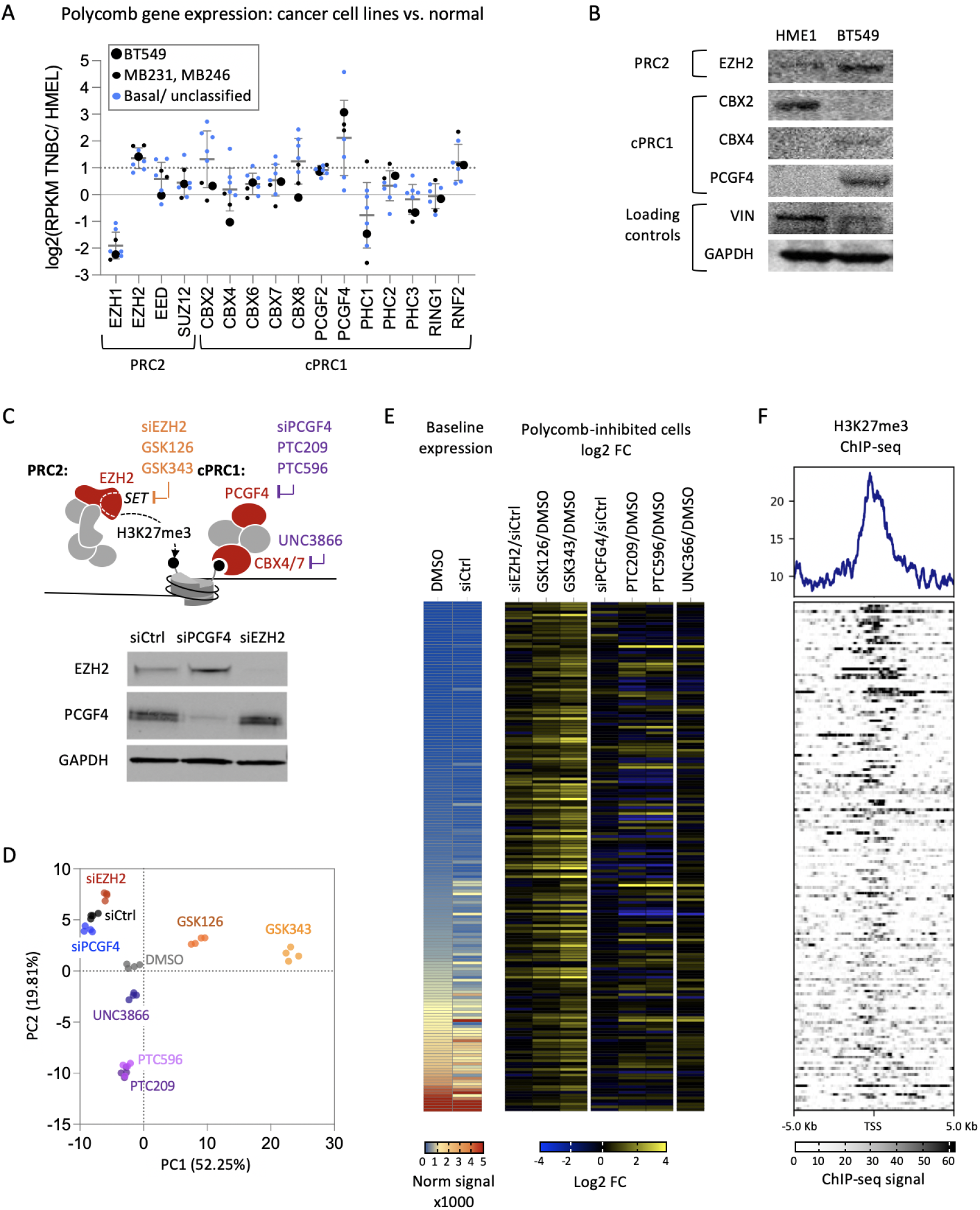
Genetic and pharmacological disruption of polycomb proteins in BT-549 cells. (A) Expression levels for polycomb subunit-encoding genes (publicly available RNA-seq data from cBioPortal) is shown for TNBC-derived cell lines versus non-cancer mammary epithelial cells (HMEL). Mesenchymal: BT-549, MDA-MB-231, MDA-MB-436; basal: HCC2157, MDA-MB-468, HCC70, HCC1806; unclassified: BT-20, HCC1500. (B) Western blots of lysates from hTERT-HME1 (non-cancer) and BT-549 cells with antibodies against the indicated polycomb proteins or loading controls. (C) Overview of polycomb targeting strategies that were tested in BT-549 for comparison in this study. Genetic knock-down via siRNA against EZH2 and PCGF4 (BMI1) in BT-549 cells was confirmed by western blot. (D) PCA plot of normalized expression levels for 177 genes in all conditions (four independent treatments each) determined by a Nanostring assay. (E) Average baseline normalized expression of 177 genes (rows) for the control samples (DMSO and siCtrl) and log2 fold change expression values for all treatments. Genes symbols and numerical values are provided in **Supplemental Table S1**. (F) H3K27me3 ChIP-seq data for BT-549 (DMSO-treated) from Lehmman et al.^19^ is shown in the transcription start site (TSS) plot, with rows corresponding to the order of rows in panel E.

We transfected BT-549 cells with siRNAs against EZH2 and PCGF4, or a control siRNA for comparison. Western blots confirmed knock-down of each protein (**Fig. 2C**). For pharmacological inhibition, we treated BT-549 for three days with IC50 concentrations (**Supplemental Fig. S2**) of EZH2 inhibitors GSK126 (IC50 5.7 μM) or GSK343 (IC50 9.5 μM), BMI1 inhibitors PTC209 (IC50 4.3 μM) or PTC596 (IC50 20.8 μM), or CBX4/7 inhibitor UNC3866 (30 μM). We analyzed total mRNA from these samples with a Nanostring assay consisting of probes for 177 genes that included low-expressed tumor suppressor genes, high-expressed cancer-promoting genes, plus other genes for broader coverage across the genome. Principal component analysis showed tight clustering of four replicates within each treatment group (**Fig. 2D**), suggesting strong reproducibility. Treatment groups showed very little overlap except for the two PCGF4 inhibitors PTC209 and PTC596.

Cells treated with EZH2 inhibitors showed the highest numbers of ≥ 2-fold upregulated genes: 88 for GSK343 and 35 for GSK126 (**Fig. 2E**). EZH2 knockdown was not entirely consistent with EZH2 inhibition (EZH2i). Ten upregulated genes were also upregulated in the EZH2i-treated groups. Compared to EZH2i, EZH2 knockdown resulted in far fewer ≥ 2-fold upregulated genes (13 total), and more ≥ 2-fold downregulated genes (4 total) (**Supplemental Table S1**). Contrary to the expectation that EZH2 inhibition would specifically affect silenced genes, both lowly and highly expressed genes became upregulated. Upregulated genes also included those with both high and low levels of H3K27me3 (**Fig. 2F**), the modification generated by EZH2. This result suggests that inhibitors of EZH2 may induce gene activation through a mechanism that does not involve loss of H3K27me3 and eviction of polycomb, e.g. by stimulating EZH2’s interactions with transcriptional activation complexes^40^.

Inhibition of PRC1 subunits PCGF4 with PTC209 or PTC596, and subunits CBX4/7 with UNC3866 induced changes in gene expression in either direction, up or down-regulation, of highly and lowly-expressed genes. Differential regulation in either direction was not restricted to loci with high levels of H3K27me3. Cells treated with PRC1 inhibitors showed fewer ≥ 2-fold upregulated genes than cells treated with EZH2 inhibitors: 22 for PTC209, 27 for PTC596, and 13 for UNC3866. A total of 8 genes were commonly upregulated between the three treatments. The most noticeable difference between PRC1 inhibition versus EZH2/PRC2 inhibition was the greater number of downregulated genes for the PRC1-inhibitor treated samples: 28 for PTC209, 25 for PTC596, and 5 for UNC3866. Decreased transcription after the inhibition of a transcriptional repressor is a counter-intuitive result that has also been observed in previous studies with rhabdomyosarcoma, where PCGF4 inhibition led to significant downregulation of dozens of genes^41,42^.

Inhibition of CBX4/7 with UNC3866 showed modest effects compared to inhibition of PCGF4. For instance 20 genes were commonly ≥ 2-fold downregulated in PCGF4 inhibitor-treated cells, whereas in CBX4/7 inhibitor-treated samples 15 were slightly downregulated (1.01 - 1.8-fold) and 3 were ≥ 2-fold downregulated. CBX4 and CBX7 are not as hyper-expressed as PCGF4 in BT-549 cells (**Fig. 2A**), therefore transcriptional regulation may not be as dependent upon CBX4/7. Furthermore, CBX4/7 inhibitor UNC3866 has a relatively modest effect on cell viability (IC50 30 μM). PCFG4 knockdown, which is an experimental standard that is used to identify PRC1-silenced genes, showed very few differentially regulated genes in common with the PCGF4 inhibitor-treated groups. Overall, only one of the 177 genes tested here, *HLA-B*, was commonly upregulated by inhibition of PRC1 and PRC2. The differential response of genes to different inhibitors and known molecular off-target effects make it difficult to use loss-of-function approaches (disruption of polycomb) to identify H3K27me3-enriched genes that are poised for epigenetic reactivation. Therefore, we used the synthetic-reader actuator fusion protein SRA-PcTF to bind H3K27me3 and stimulate transcriptional activation via recruitment of Mediator subunits in polycomb-repressed chromatin.

### SRA-PcTF activates low-expressed genes near H3K27me3-adjacent enhancers

To identify transcriptionally silenced H3K27me3-enriched genes that are poised for activation, we used SRA-PcTF to stimulate the pre-initiation complex through interactions between the SRA’s C-terminal VP64 domain and endogenous MED15 and MED17^26,27^. In a previous study^29^, we transfected BT-549 cells with SRA-expressing plasmids, extracted mRNA 24, 48, and 72 hours after transfection, and analyzed these samples with RNA-seq (**Fig. 3A**). In this study, we aligned the reads and analyzed the data using STAR and DESeq2 and identified 122 protein-coding, upregulated differentially expressed genes (UpDEGs, ≥ 2-fold activation, adjusted *p* < 0.05) collectively at 24, 48, and 72 hours after SRA transfection (**Fig. 3B, C**).

**Figure 3.**
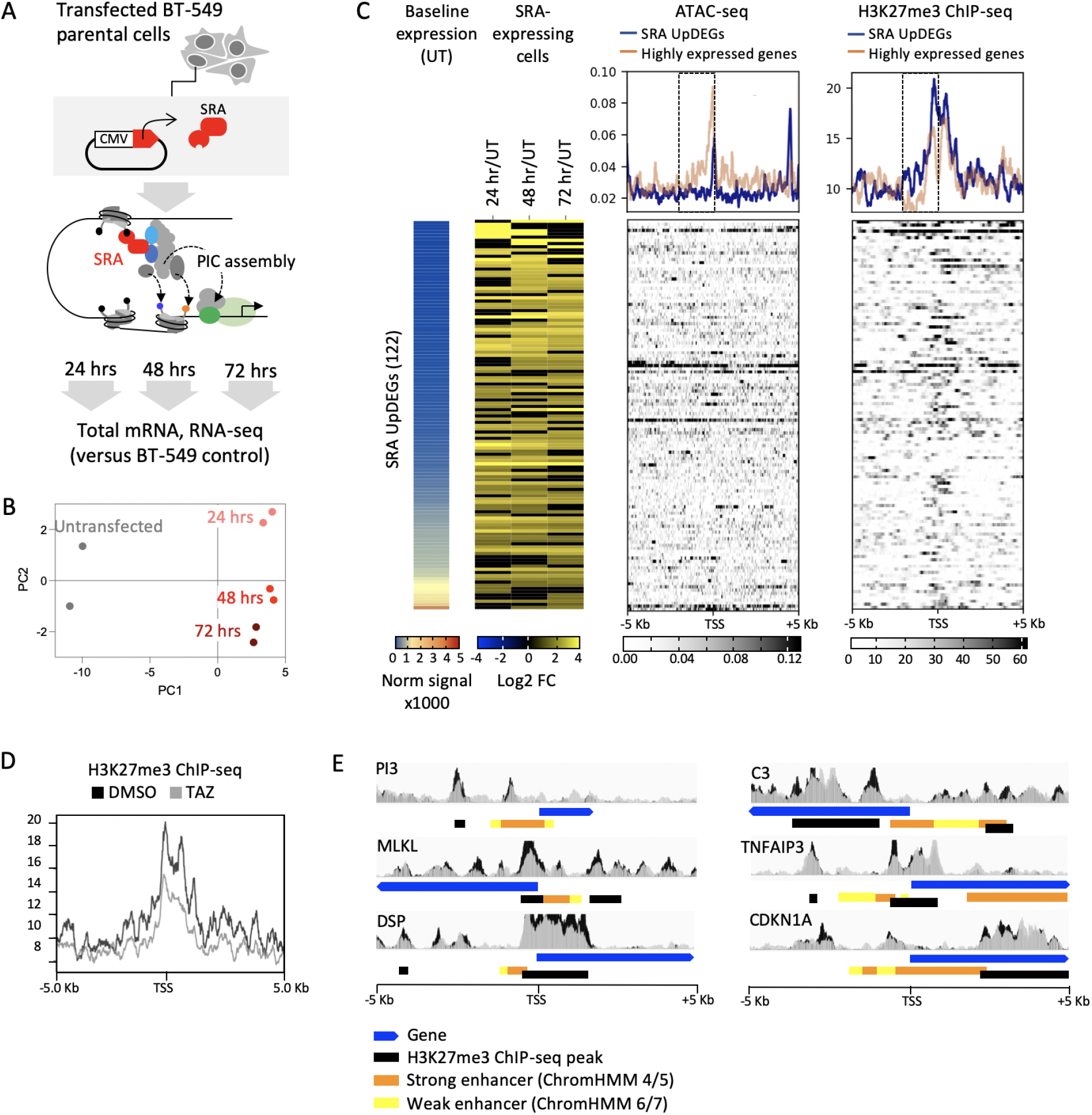
Transcription and epigenomic profiling of SRA-expressing BT-549 cells. (A) BT-549 cells were transfected with SRA-expressing plasmid DNA and analyzed with RNA-seq 24, 48, and 72 hours after transfection (2 replicates per condition). (B) Principal component analysis showed strong reproducibility of RNA-seq data from two biological replicates (independent transfections) per condition. (C) 122 SRA UpDEGs (rows) are sorted from low to high baseline expression. Log2 fold change at each time point post-transfection is shown in the color gradient map. The top 100 highly expressed genes are included for comparison. Transcription start site (TSS) plots for H3K27me3 ChIP-seq and ATAC-seq signals show mean signals across 10 Kb regions centered at the TSS’s of each group of genes. Expression values for SRA UpDEGs and the top 100 highly expressed genes are provided in **Supplemental Table S2**. (D) The TSS plot shows mean H3K27me3 ChIP-seq signals for BT-549 cells control-treated with DMSO or treated with the EZH2 inhibitor Tazemetostat. (E) Wiggle (WIG) tracks show H3K27me3 ChIP-seq data from DMSO (black) or Tazemetostat-treated (gray) BT-549 cells. H3K27me3 ChIP -seq peaks and annotations for chromatin features from the ENCODE Broad ChromHMM dataset^43^ are shown for six representative SRA UpDEGs that are located near strong enhancers.

We expected SRA UpDEGs to be located within transcriptionally repressed regions that are depleted of transcription-activating proteins and contain nucleosomes with H3K27me3. Baseline expression levels (normalized average counts) from RNA-seq data for the 122 SRA UpDEGs had a median value of 103.33, which was lower than median for the top 100 highly expressed genes (7125.16). Public RNA-seq data for the 122 SRA target genes (UpDEGs) that were identified in our RNA-seq experiment show low median levels of transcription in BT-549 and other TNBC cell lines compared to HMEL normal mammary epithelial cells (**Supplemental Fig. S3**). The low expression levels that we observed suggest chromatin-mediated epigenetic silencing.

To measure DNA-binding transcription factor occupancy, we analyzed 50-115 bp ATAC-seq fragments, which correspond to Tn5 hypersensitive sites (THSS) bound by transcription factors. We observed lower ATAC-seq signals in regions up to 2 Kb upstream of the transcription start sites (TSS’s) of the SRA UpDEG group compared to the top 100 highly expressed genes, suggesting fewer THSS’s and lower transcription factor occupancy. Next, we analyzed H3K27me3 ChIP-seq signals^19^ as an indicator of polycomb chromatin formation. We observed higher H3K27me3 ChIP-seq signals in the same 2 Kb regions (**Fig. 3C**). H3K27me3 signals were decreased in cells treated with the EZH2 inhibitor Tazemetostat, indicating that histone K27 methylation at these sites is dependent upon EZH2 activity (**Fig. 3D**). Our previous work with U2-OS, K562, and SK-N-SH cell lines suggested that SRA-PcTF activates genes for which H3K27me3 is located near enhancer regions^28^. To determine the location of H3K27me3 at breast epithelial cell enhancers, we investigated annotated enhancer regions using chromatin state segmentation data from ENCODE for the human mammary epithelial cell line HMEC^43^. For genes with strong enhancers located within 10 kb around the transcription start site, we observed that EZH2-dependent H3K27me3 peaks lie within or near enhancers located upstream or surrounding the transcription start sites of SRA UpDEGs (**Fig. 3E**).

### H3K27me3-targeting by the SRA-PcTF transcriptional regulator activates tumor suppressor and cancer promoting genes in triple negative breast cancer cell line BT-549

The vast majority of differentially expressed genes in SRA-expressing cells were upregulated (**Fig. 4A**). Fifty-two (42%) of the UpDEGs were shared across all replicates and time points, which represent six different samples (**Fig. 4B**). The SRA UpDEGs include genes involved in cell death (necroptosis and apoptosis), cell cycle arrest, inhibition of invasion, and immune surveillance (**Fig. 4C, Supplemental Table S3**). UpDEGs that positively modulate necroptosis are part of the TNF signaling pathway (KEGG hsa04668, hsa04217). *STAT1* and *STAT2* transcription factors activate the expression of core members of the necrosome^44^, and *TLR3* promotes necrosome formation^45^. *MLKL* is stimulated by the necrosome, promotes sodium influx, cell membrane rupture, and inflammation by the release of cytokines^46^ such as *IL1A*, *IL1B* and *CXCL10* which are also upregulated by the SRA. *PMAIP1* promotes apoptosis (KEGG hsa04210) through the release of apoptogenic proteins from the mitochondria^47^. UpDEGs that support cell cycle arrest include members of the cellular senescence pathway (KEGG hsa04218). *CDKN1A* blocks cell cycle progression by binding to and inhibiting cyclin-dependent kinases^48^. *HLA-A*, *HLA-B*, *HLA-C*, and *HLA-E* act upstream of *CDKN1A* expression through DNA damage signaling, and *IL1A* supports the senescence-associated secretory phenotype. The UpDEGs involved in cellular migration include genes that mediate ECM-receptor interactions (KEGG hsa04512) and tight junctions (KEGG hsa04530). *ITGA2* and *CLDN1* are repressed in aggressive breast cancer^49,50^, although some studies suggest that *ITGA2* promotes breast cancer metastasis^51^. Immune surveillance pathways are the most represented amongst the UpDEGs, and these genes are known to support the immunosuppression of cancer. We observed significant downregulation of only four genes including *FGFR2*, *CCN2*, *GFRA1*, and *GAS1*. We expect the VP64-containing SRA protein is only capable of inducing activation, therefore downregulation of these genes could be a consequence of increased expression of a transcriptional repressor protein or microRNA. We could not identify potential negative regulators in the SRA UpDEG subset.

**Figure 4.**
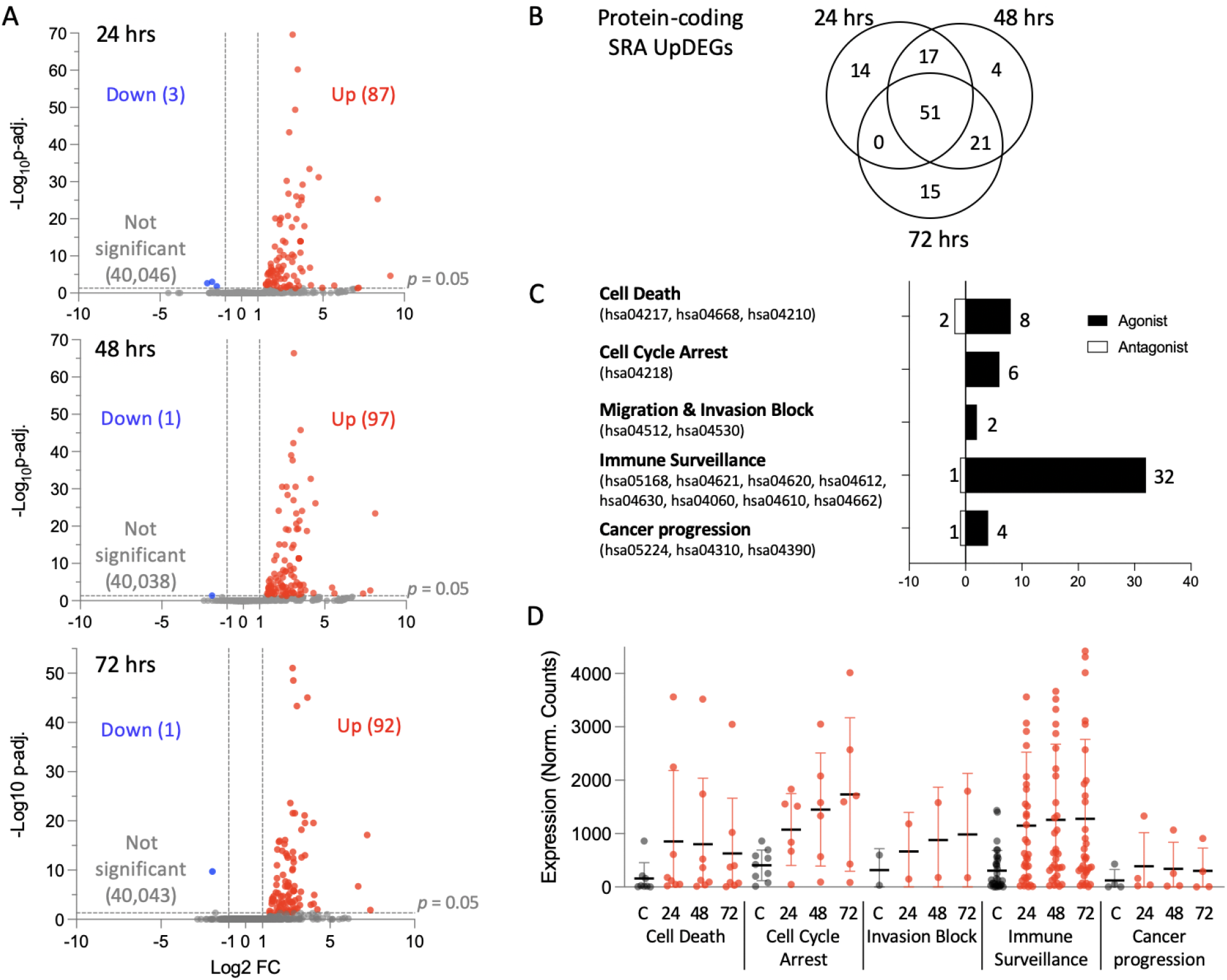
Transcription profiling of SRA-PcTF-expressing BT-549 cells. (A) Volcano plot of log2 fold change ratios for each time point. Differentially regulated genes (DEGs, |FC| ≥ 2, adjusted p < 0.05) are highlighted blue for downregulation or red for upregulation. (B) Venn diagram of protein-coding SRA-upregulated DEGs (UpDEGs) from each time point. (C) Functional categorization of SRA UpDEGs. KEGG pathway IDs are shown. Genes are listed in Supplemental Table 3. (D) Expression values (average normalized counts for two replicates) of UpDEGs within the five functional categories in panel C. Data for the Cell Death group exclude antagonists *BIRC3* and *TNFAIP3*. Data for the Cancer Progression group exclude the antagonist *CDKN1A*.

Some cancer-promoting genes were also upregulated by the SRA. Five UpDEGs including *BRC3*, *BMP2*, *DLL1*, and *LEF1* are members of cancer progression pathways that involve Wnt signaling and Hippo signaling (KEGG hsa05224, hsa04310, hsa04390). *BIRC3* and *TNFAIP3* negatively modulate apoptosis and necroptosis through the TNF signaling pathway (KEGG hsa04668). The activation of genes that either inhibit or promote cancer cell proliferation and survival led us to ask whether one set might have a dominant impact on cell phenotype. We grouped the genes into categories based on their associated pathways and observed that in SRA-expressing cells median expression values for the anti-cancer genes are higher than median expression values for the cancer-promoting genes (**Fig. 3E**). The SRA-induced expression profile, which shows higher expression of pathways associated with cell death, cell cycle arrest, and inhibition of migration and invasion compared to cancer-promoting genes, prompted us to determine changes in proliferation and invasion of cells into an extracellular matrix *in vitro*.

### SRA-PcTF expression reduces BT-549 spheroid size in a Matrigel dependent manner

Others have observed that inhibitors of EZH2 (e.g. GSK126) and BMI1 (e.g. PTC209) disrupt polycomb-mediated epigenetic silencing, and block TNBC aggression in tumor xenograft models. Presumably, tumorigenic processes such as proliferation and invasion are halted when polycomb complexes are disabled and tumor suppressor genes become transcriptionally active. Therefore, we surmised that recruitment of transcriptional activators to polycomb-repressed promoters via a synthetic reader-actuator (SRA) would activate tumor suppressor genes and block growth and invasion. To test this idea, we measured growth inhibition in a 3-D spheroid model (**Fig. 5A**).

**Figure 5.**
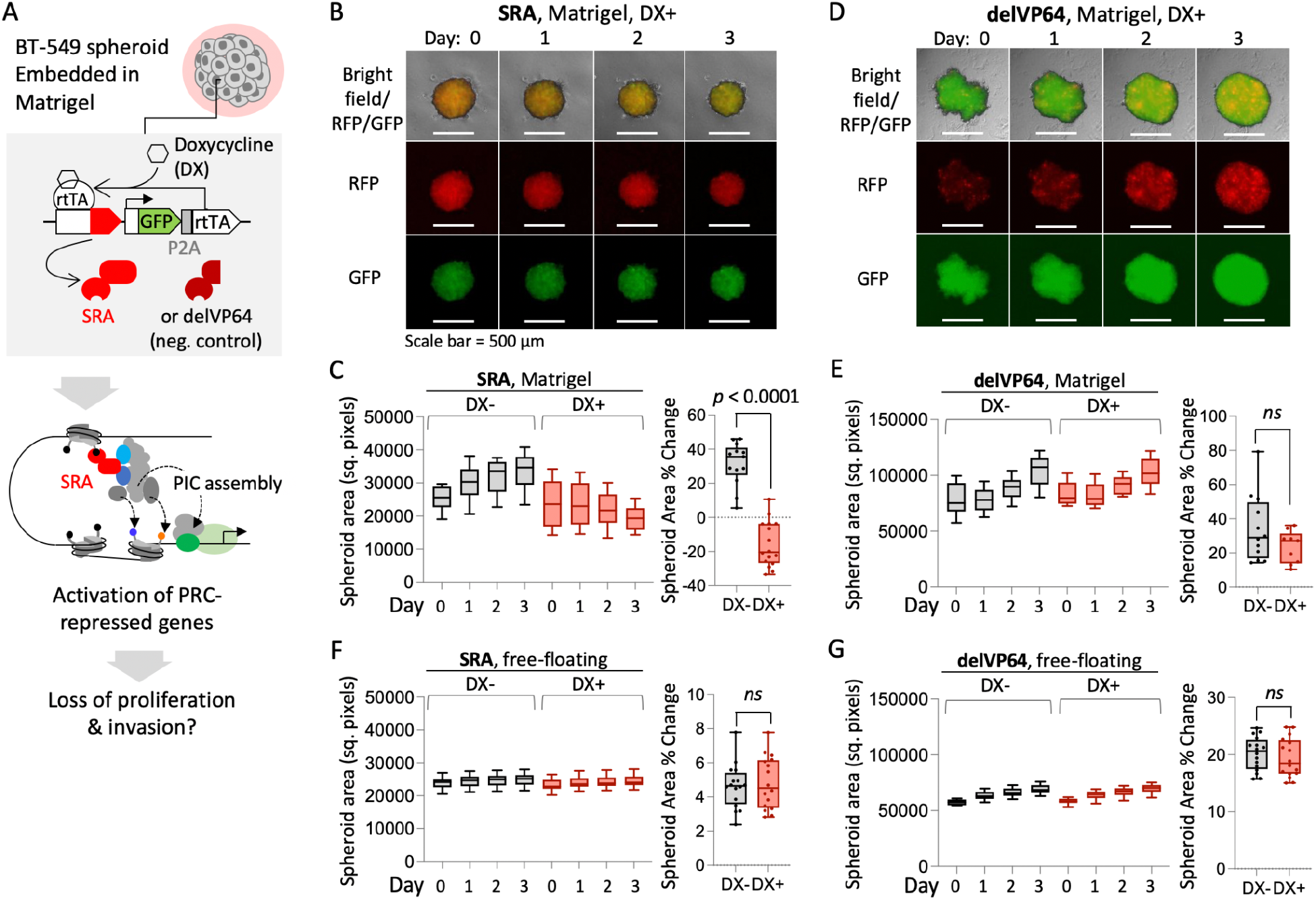
(A) Experiment overview: activation of transgenic SRA expression with doxycycline (DX). (B) Representative spheroid image showing reduction in size of SRA-expressing BT-549 spheroids up to 3 days after embedding. (C) The left box plot shows areas of SRA-expressing, Matrigel-embedded spheroids without (*n* = 13) or with DX (*n* = 16) over time. The right box plot shows percent changes in areas for individual spheroids at day 0 versus day 3. (D) Deletion of the SRA VP64 domain rescues spheroid growth. (E) Spheroid areas and percent change are shown for delVP64-expressing spheroids without (*n* = 12) or with DX (*n* = 9). (F, G) The same analyses were performed for free-floating SRA-expressing and delVP64-expressing spheroids without or with DX (*n* = 16 for each group). *p* values: two-tailed, unpaired t-test. Ns = not significant, *p* >0.1.

To enable SRA expression throughout the spheroid, we engineered BT-549 cells that carry an SRA transgene (pSBtet-GP system^52^) under tight control of doxycycline (DX) (**Fig. 5A**). Harvested cells were allowed to form spheroids in low-binding u-bottom 96-well plates (1000 cells per well) in liquid media for three days. 72 hours after seeding, Doxycycline was added to the spheroids to induce expression of pSBtet-GP. Spheroids were embedded in 80-90% Matrigel without or with 1 μg/μL doxycycline, then imaged the same day (day 0) and over the next three days (days 1 - 3). SRA-expressing spheroids showed a reduction in size (−20% median spheroid area) while spheroids without activation of the pSBtet-GP promoter showed an increase in size (+35% median spheroid area) (**Fig. 5B, C**). Spheroids carrying a truncated SRA (deletion of the VP64 transcriptional activator domain) showed similar increases in growth without or with activation of pSBtet-GP (+35% media spheroid area) (**Fig. 5D, E**).

The reduction in spheroid size observed for SRA DX+ might be caused by cell death within the spheroid body or the death of cells that have migrated away from the spheroid. To determine if reduced spheroid size occurs within the spheroid body, independently of growth or invasion in the Matrigel, we seeded spheroids for three days, continued to culture the spheroids in liquid media without or with 1 μg/μL doxycycline, and imaged the spheroids on days 0 - 3 as we did for embedded spheroids. Expression of the SRA resulted in no change in spheroid size (**Fig. 5F, G**). Overall these results suggest that activation of the pre-initiation complex via SRAs at polycomb-repressed genes induces an anti-proliferative state, which may specifically affect invasive spheroids.

### SRA-PcTF expression stimulates an anti-invasive and apoptotic state in 3-D spheroids

SRA-expressing cells showed increased expression of *ITGA2* and *CLDN1* in our RNA-seq experiment (**Fig. 6A**), and these genes mediate cell migration and invasion. Others have reported that artificial overexpression of *ITGA2* or *CLDN1* blocks breast cell migration *in vitro^49,50^*, suggesting that increased expression of these genes may be sufficient to halt spheroid invasion. Additional SRA UpDEGs such as *DSP* (desmoplakin), *TFPI2*, and *MME* (neprilysin) may also inhibit migration^53–55^. *CCN2*, which is positively associated with breast cancer invasion^56^, becomes downregulated in SRA-expressing cells. To determine the effect of SRA-mediated gene regulation on invasion, we established an invasion model using spheroids embedded in Matrigel. Both engineered cell lines carrying either the SRA- and delVP64-encoding transgene (see **Fig. 5**) showed the capacity to develop an invasive margin of cells 3 days after embedding (**Fig. 6B**).

**Figure 6.**
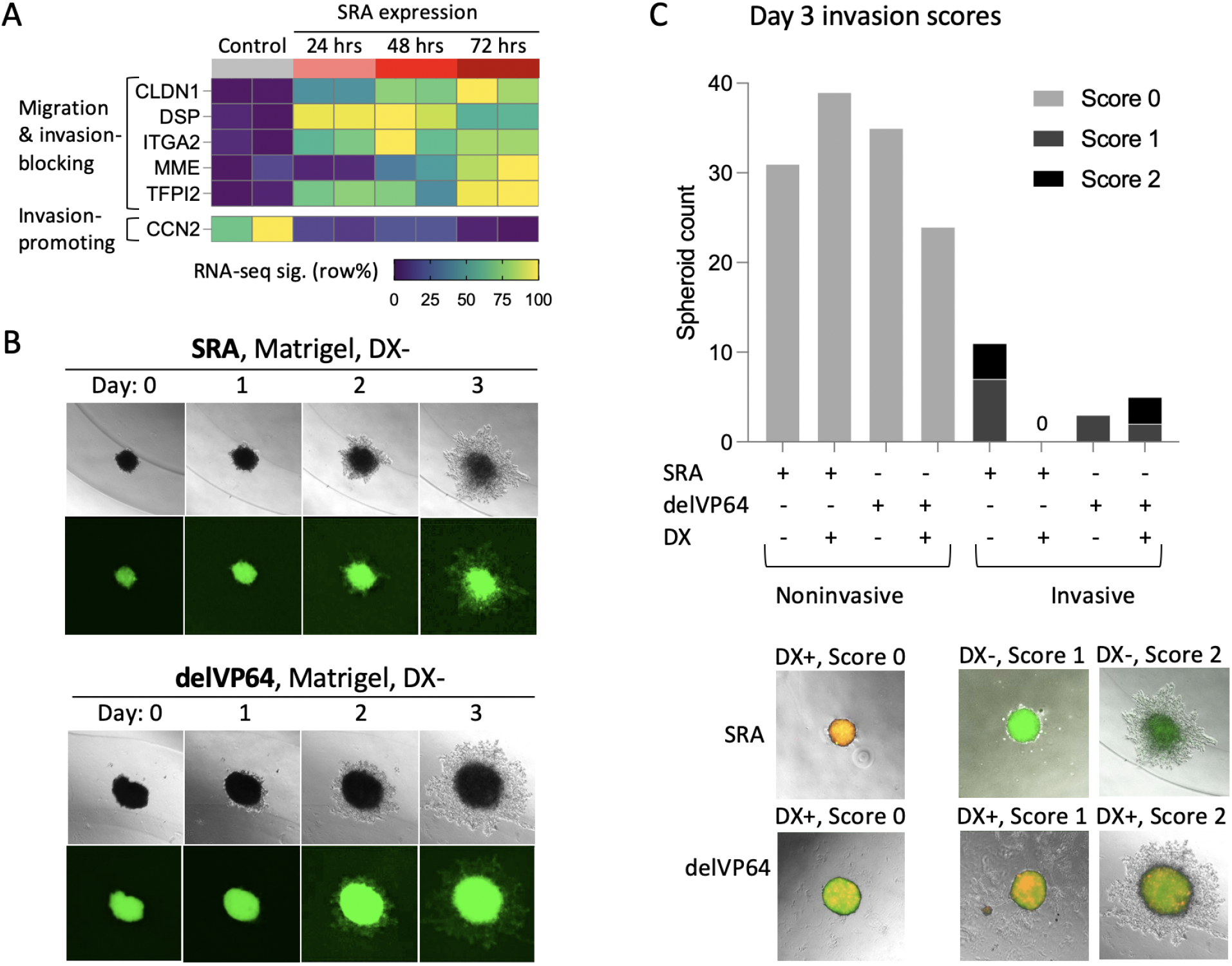
Measurement of the invasive capacity of 3-D spheroids expressing SRAs or the control delVP64 protein. (A) Normalized expression values for SRA UpDEGs involved in migration and invasion (RNA-seq data from **Fig. 3**). (B) Representative images of transgenic BT-549 spheroids (without doxycycline) that undergo invasion three days after being embedded in Matrigel. (B) Embedded spheroids were scored for invasion after 3 days of growth. Representative images are shown for each invasion score.

We compared invasion frequency in SRA-expressing spheroids versus non-expressing spheroids, and spheroids that expressed the delVP64 negative control protein. After inducing SRA or delVP64 expression with doxycycline, spheroids were embedded in Matrigel and observed. Spheroids were given a score based on the degree of invasion observed. Spheroids with no invasion, a small margin of invading cells, or a large invasive margin were given a score of 0, 1, or 2 respectively (**Fig. 6C)**. Three days after spheroids were embedded, we detected 11 invasive events of 42 non-expressing (DX-) spheroids. No invasion events occurred for 39 SRA-expressing (DX+) spheroids (**Fig. 6C**). DelVP64 negative control spheroids, which lack the VP64 domain that recruits transcriptional activating proteins MED17 and MED25, showed invasion events in both the mCherry-expressing (DX+) and non-expressing (DX-) spheroids. The delVP64 negative control had 5 invasion events out of 29 DX+ spheroids. In the non-expressing (DX-) delVP64 spheroids, 3 invasion events occurred for 38 spheroids embedded (**Fig. 6C**). These results suggest that SRA UpDEGs that are members of invasion-blocking pathways (i.e. hsa04512: ECM-receptor interaction, hsa04530: tight junction), are activated at sufficient levels to halt invasion.

To determine if apoptosis also occurs during SRA expression, we performed staining of annexin V in SRA-expressing cells. We used transient transfections of SRAs instead of the stable lines because the transgenic chromosomal GFP marker is incompatible with annexin V (FITC channel). To test the hypothesis that PCD-mediated binding is required for tumor suppressor activation and apoptosis, we built a set of SRAs that included two wildtype H3K27me3-binding domains, or a mutation (F11A) in the aromatic binding pocket of one or both PCDs (**Fig. 7A**). Three days after lipofectamine transfection, we detected 4-5% RFP-positive cells (of ~85,000 total). 55% of RFP-positive SRA-expressing cells were annexin V-positive, which was higher than the single mutant (33%) and double mutant (25%). RFP-positive SRA cells also showed a rounded morphology similar to that of paclitaxel-treated cells (10 nM), measured as smaller area per cell (**Fig. 7B**). Therefore, Mediator recruitment to polycomb-repressed sites by SRAs enhances tumor suppressor transcription, stimulates apoptosis, and blocks proliferation and invasion of TNBC cells.

**Figure 7.**
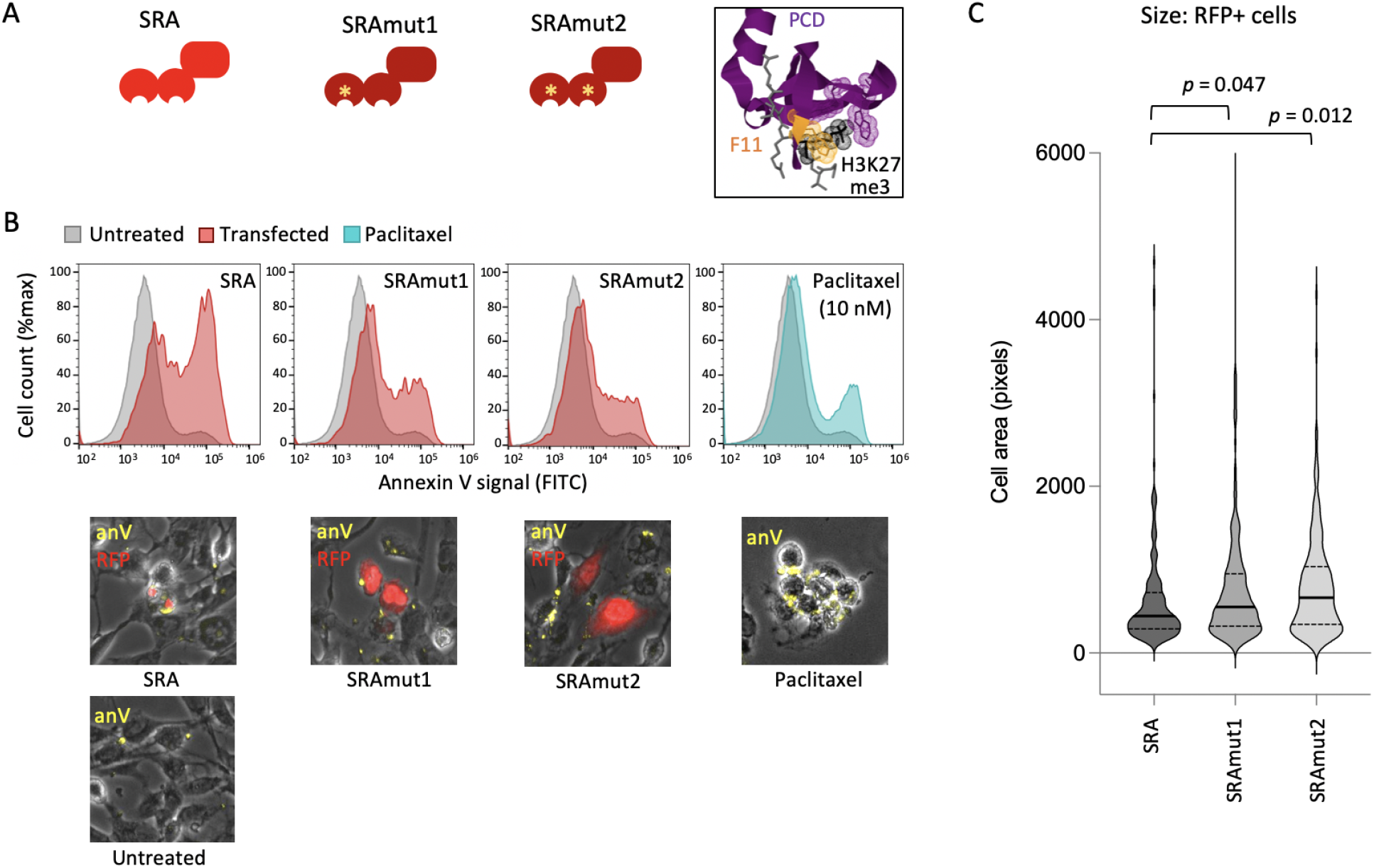
Apoptosis and cell size reduction in SRA-expressing cells. (A) Schematic of SRA variants carrying two wild type H3K27me3-binding domains (PCD), or an F11A mutation. The PCD 3-D structure is from the Protein Data Bank model PDB 3I91^57^. (B) Annexin V (anV) staining and flow cytometry of BT-549 cells expressing RFP-tagged SRA proteins. Point mutations in the H3K27me3-binding PCD domains (F11A) rescue viability. Paclitaxel-treated cells (10 nM) are a control for apoptosis. P.I. staining was omitted because it is incompatible with detection of RFP from the SRA protein. (B) SRA-expressing cells show reduced size compared to cells that express mutated SRAs. Areas of 200 RFP-positive cells per condition were measured by microscopy and BioTek Gen5 software. *p* values: two-tailed, unpaired t-test.

## DISCUSSION AND CONCLUSIONS

We used an engineered chromatin protein to identify chromatin-repressed genes that are capable of having their transcription restored in triple negative breast cancer cells with high levels of polycomb expression. Our results demonstrate how synthetic reader-actuators can be used as an alternative to polycomb-disrupting agents, such as small molecule inhibitors and genetic knockdown, which show inconsistent effects on transcriptional activation. Removal of polycomb repressive complexes (PRCs) from DNA by pharmacological inhibition or genetic knockdown is expected to allow assembly of the preinitiation complex (PIC) at promoters. However, with this approach there is no step to actively recruit or restore transcriptional activators. Studies using mouse embryonic stem cells showed that depletion of either PRC1 or PRC2 did not increase DNA accessibility at polycomb-occupied genes^17,18^ which suggests incomplete conversion to a transcriptionally active state. Polycomb eviction via epigenetic inhibitors activates immune signaling genes in triple negative breast cancer cells, and to a lesser extent apoptotic genes. TAZVERIK (tazemetostat) inhibits the SET domain (histone methyltransferase) of EZH2, and is FDA approved for follicular lymphoma and metastatic sarcoma^58^. However, tazemetostat only reduces TNBC cell viability by 50%, even at high concentrations (20 μM)^19^ raising the possibility that polycomb inhibition does not fully activate apoptotic gene expression in TNBC. Inhibition of EZH2 with GSK126 suppresses TNBC metastasis in patient-derived and MDA-MB-231 xenograft tumors in mice but does not significantly reduce the size of the primary tumor^16,59^. In certain contexts, EZH2 inhibition may even be anti-therapeutic. In hypoxic TNBC cells (MDA-MB-231) GSK126 stimulates EZH2/FOXM1 complex formation, transcriptional activation of matrix metalloprotease enzyme genes, and cell invasion in transwell assays^40^.

When we tried to identify activatable H3K27me3-enriched genes in BT-549 cells by inhibiting specific polycomb complex subunits, both lowly and highly expressed genes became up- or down-regulated, and this effect was not limited to H3K27me3-enriched genes (**Fig. 2**). Inhibitors of EZH2 might induce gene activation through a mechanism that does not involve loss of H3K27me3 and eviction of polycomb, e.g. by stimulating EZH2’s interactions with transcriptional activation complexes in certain contexts such as Wnt and estrogen signaling^60^ and nuclear factor-kappa B gene regulation^61^. Polycomb inhibition can also lead to the loss of transcription, as we observed with our Nanostring profiling assay with BT-549 TNBC cells (**Fig. 2**). A ChIP-seq study in TNBC cells showed that PRC1 complexes (containing CBX8, MEL18/PCGF2, RING1B/RNF2, and PHC2) accumulated at enhancers that were devoid of H3K27me3 and enriched with the transcriptional activator BRD4^62^, suggesting that PRC1 might maintain active expression of some genes. In our previous studies with rhabdomyosarcoma cell lines, PCGF4 inhibition lead to significant downregulation of dozens of genes^41,42^ possibly through elevated expression of kinase encoding genes, phosphorylation-mediated inactivation of YAP/TAZ/TEAD, and silencing of YAP/TAZ-target promoters. In another group’s study, treatment of glioblastoma multiforme (GBM) cells with PTC596 affected protein levels of transcriptional regulators EZH2, FOXG1, and SOX2. PRC2 genes were de-repressed as expected, but a different subset of genes regulated by SOX2 were downregulated. Off-target effects of polycomb loss-of-function approaches make it difficult to investigate the regulation of H3K27me3-enriched genes.

When we expressed SRA-PcTF in BT-549, the vast majority of differentially regulated genes were activated (UpDEGs). The SRA UpDEGs included low-expressed genes that were located near strong enhancers adjacent to H3K27me3 peaks (**Fig. 3**). These enhancers showed a depletion of Tn5 hypersensitive sites (ATAC-seq) compared to the top highly expressed genes, suggesting lower transcription factor occupancy. These results provide insights into the state of epigenetically silenced genes in TNBC and how their expression can be restored. Transcription-initiating proteins might be displaced by the occupancy of hyper-expressed polycomb proteins. Alternatively, the loss of key transcriptional regulators such as TP53^19,21,22^ and components of BAF SWI/SNF^19,24^ might permit the accumulation of polycomb and H3K27me3 at enhancers. In either case, the introduction of a protein that binds H3K27me3 recruits parts of the core transcriptional machinery, including MED17 and MED25, through interactions with VP64 (**Fig. 1C**) is sufficient to restore transcription.

A large body of evidence suggests that polycomb hyper activity confers an advantage to cancer cell proliferation and survival by silencing tumor-suppressive genes. However it is not yet fully understood whether repression is restricted to tumor suppressors since polycomb complexes are often distributed across dozens of genes. The results from our study with SRA-PcTF suggests that activatable genes in polycomb-repressed chromatin include both tumor suppressive and cancer-promoting activities. Therefore nonspecific activation of these genes might stimulate competing pathways. We observed that the majority of the SRA UpDEGs include tumor suppressor genes involved in cell death (necroptosis and apoptosis), cell cycle arrest, inhibition of migration and invasion, and immune surveillance (**Fig. 4**), and these genes showed greater transcriptional activation than the small fraction of cancer-promoting SRA UpDEGs. Transcriptional reprogramming via SRA-PcTF was accompanied by anti-cancer cellular phenotypes including a reduction of spheroid size, loss of extracellular matrix invasion, and increased apoptosis (**Fig. 5, 6, 7**). These results suggest that chromatin-targeted recruitment of endogenous transcription factors to polycomb-repressed sites effectively restores tumor suppressor gene expression in TNBC cells. Anti-proliferative and anti-invasive epigenetic reprogramming were observed in the absence of the off-target effects that are associated with inhibition of EZH2 and PCGF4. These results demonstrate the utility of synthetic reader-actuators for investigating the therapeutic potential of direct transcriptional reprogramming of coregulated genes in TNBC.

## METHODS

### 2-D cell culture

All cell lines were cultured at 37°C and 5% CO_2_. hTERT-HME1 (ATCC CRL-4010) cells were cultured in MEGM media (Lonza, CC-3150) without GA-1000. BT-549 (ATCC HTB-122) cells were cultured in RPMI-1640 media (Gibco, 11875119) supplemented with 0.0008 mg/mL human insulin. RPMI-1640 media contained 10% fetal bovine serum (Atlanta Biologicals, S10350) and 1% penicillin/streptomycin solution (Gibco, 15140122).

### Western blot

BT-549 and hTERT-HME1 cells were grown to ~80% confluency in 10 cm tissue culture-treated plates and harvested with 0.25% trypsin-EDTA (Thermo Fisher Scientific, 25200056). Lysates were extracted using RIPA lysis buffer and insoluble protein was cleared by centrifugation at 16,000xg for 30 min. Lysates were electrophoresed in a 12% NuPAGE Bis Tris Protein gel (Fisher Scientific, NP0336BOX) and transferred to nitrocellulose membranes (Bio-Rad, 1704158). Primary staining was performed on separate blots with the following antibody dilutions in 5% BSA/TBST: rabbit anti-EZH2 1:1000 (Cell Signaling Technology, 5246S), rabbit anti-CBX2 1:500 (Bethyl Laboratories, A302-524A-T), rabbit anti-CBX4 1:1000 (Abcam, ab242149), rabbit anti- PCGF4/ BMI1 1:1000 (Cell Signaling Technology, 6964S), rabbit anti-VIN 1:1000 (Cell Signaling Technology, 13901S), rabbit anti-GAPDH 1:1000 (Cell Signaling Technology, 2118S). Secondary staining was performed with 1:30000 goat anti-rabbit IRDye 800CW-conjugated (LI-COR BioScience, 926-32211) in 5% BSA/TBST. Post primary and secondary stain washes were done four times with TBST. Signal detection was performed using an iBright imager (Thermo Fisher, A44241).

### Polycomb protein knockdown with siRNA

BT-549 cells were seeded at 2.0E5 cells per well in 6-well plates. After 24 hours, BT-549 cells were transfected using DharmaFECT 1 (Horizon Discovery, T-2001-03) and 25 nM of an siRNA ON-TARGETplus SMARTpool (Horizon Discovery, BMI1: L-005230-01-0010, EZH2: L-004218-00-0010) or ON-TARGETplus Nontargeting Control Pool (Horizon Discovery, D-001810-10-20). Cells were harvested for total RNA (RNeasy Mini Kit - Qiagen, 74104) and protein 72 hours after transfection. Western blots were performed as described above.

### Nanostring assay

96-well plates were coated with 2% ECM-supplemented medium (Millipore Sigma, E0282) which was allowed to solidify for 4 hours. Cells were seeded at approximately 1.0E4 per well. Drugs were brought to 10 mM in DMSO. 24 hours after seeding, 50 μL of diluted compound was added to each cell sample to generate final concentrations of EZH2 inhibitors GSK126 (5.7 μM; Sigma, 5005800001) or GSK343 (9.5 μM; Sigma, SML0766), BMI1 inhibitors PTC209 (4.3 μM; Selleck Chemicals, S7372) or PTC596 (20.8 μM; Selleck Chemicals S8820), and CBX4/7 inhibitor UNC3866 (30 μM; Sigma, SML2408). Cells were harvested for RNA at 72 hours after application of inhibitors. RNA was isolated using a RNeasy Micro Kit (Qiagen, 74004). The Nanostring assay was performed at the Emory Integrated Genomics Core (EIGC) using a custom codeset that included 177 protein-coding genes and three housekeeping control genes (ACTB, CHMP2A, GAPDH).

### RNA-seq of transiently transfected cells

BT-549 cells were cultured, transfected, and processed at 24, 48, and 72 hours after transfection (or no transfection for the negative control) to collect total RNA as we have described previously^29^. Raw reads (GSM2773005-12) were trimmed with Trimmomatic and aligned via seed searching using STAR with the most recent human genome (GRCh38/hg38). BAM files were sorted and marked with picard and differential gene expression analysis was performed with DESeq2. DESeq2 analysis uses a negative binomial distribution algorithm to recognize genes with a p value ≤ 0.05 and log2 fold change difference of ≤ −1 or ≥ 1.

### ATAC-seq

Roughly 1.0E6 BT-549 cells were harvested from 6-well plates and frozen in 50% RPMI, 40% FBS, 10% DMSO. Sample processing, library preparation, and sequencing was performed by Novogene. All libraries were sequenced using an Illumina NovaSeq 6000 instrument in a 50 bp paired-end format. Paired reads were aligned to the human hg38 reference genome using Bowtie2^63^ with default parameters, except −X 2000. PCR duplicates were removed using Picard Tools^64^. To adjust for fragment size, we aligned all reads as plus (+) strands offset by +4 bp and minus (−) strands offset by −5 bp^65^. Reads were separated by size into 50-115 bp fragments, which correspond to Tn5 hypersensitive sites (THSS) bound by transcription factors (TFs), and 180-247 bp fragments, which correspond to mononucleosomes^66–68^. Peaks were called in the THSS and mono-nucleosome fractions using MACS2 with default parameters^69^. ATAC-seq peaks that overlap with the ENCODE blacklist for hg38 were excluded from further analysis^70^. TSS plots were generated by DeepTools computeMatrix and plotHeatmap using bigwig files for THSS signals.

### ChIP-seq data analysis

Wiggle and BED files were visualized with the Integrative Genomics Viewer (IGV 2.8.2)^71^ using the GRCh37/hg19 human genome reference. Bigwig files for BT-549 DMSO and Tazemetostat, and the bed file for DMSO versus Tazemetostat peaks were provided by B. Lehmann. Annotated ChromHMM features for HMEC cells were visualized in IGV using the GRCh37/hg19 reference sequence and the BED file wgEncodeEH000786 from UCSC: Chromatin State Segmentation by HMM from ENCODE/Broad.

### SRA-expressing stable transgenic cell lines

The BT549-DBN021 (SRA) and BT549-DBN025 (delVP64) cell lines were generated by transient transfection with pSBtet-GP plasmid and Lipofectamine complexes, followed by selection with puromycin as described previously^29^. Monoclonal isolates were reselected by limiting dilution in media with 0.5 μg/μL puromycin to 1 cell per 100 μL in 96-well plates. Plates were examined for wells that had 1-2 cells per well. These isolated cells were allowed to proliferate to 80-100% confluency, expanded to 12-well plates, sub-sampled and tested for dox-inducible uniform RFP expression, and then expanded to 10 cm dishes before cryogenic storage in 10% DMSO in FBS.

### 3-D spheroid culture

3-D spheroid cultures were seeded at 1.0 - 1.5E3 cells per well in a total volume of 200 μL of growth media in ultra low-binding round bottom 96-well plates (Corning, 7007). After 3 days, wells were inspected for round clusters. Doxycycline (1 μg/μL) was applied to the spheroids 3 days after seeding. For invasion experiments, spheroids were embedded in a basement membrane matrix, Matrigel (Corning, 356234). Briefly, solidified beds of Matrigel were prepared in a 6-well plate with four 25 μL beds of Matrigel per well. The Matrigel beds were allowed to solidify for 30 minutes at 37°C. Individual spheroids were transferred to microtubes and the culture media was removed. 20 μL Matrigel was added to the spheroid in each microtube, and the spheroid plus Matrigel was carefully transferred onto a solidified bed of Matrigel. The embedded spheroids were allowed to solidify at room temperature for 5 minutes, then at 37°C for 30 minutes. 4 mL culture media was added to each well. Doxycycline was added to Matrigel and culture media (final concentration 1 μg/mL) for the spheroids that received an earlier treatment of doxycycline. Embedded spheroids were imaged using an EVOS M5000 Cell Imaging System (ThermoFisher Scientific, AMF5000). Images were taken on the initial day of embedding and 3 additional days post-embedding.

### Annexin V staining and flow cytometry

SRA constructs (SRA: SA0020_MV1, SRAmut1: SA0021_MV1, SRAmut2: SA0022_MV1) were generated via Golden Gate assembly^72^ and cloned into the XbaI and SpeI sites of expression vector MV1. Annotated sequences are available at Benchling (https://benchling.com/hayneslab/f_/rmSYkAAU-synthetic-chromatin-actuators-2-0/). BT-549 cells were plated at 2.0E5 cells per well in 6-well plates in 2 mL complete media, grown overnight, and transfected with Lipofectamine LTX complexes: 1 μg plasmid DNA, 5.0 μL Lipofectamine LTX (Thermo Fisher, 15338030), and 2.5 μL PLUS reagent brought to a final volume of 500 μL with OptiMEM (Thermo Fisher, 11058021). Additional untransfected samples were treated with paclitaxel at a final concentration of 10 nM. After 72 hours, transfected cells were collected for Annexin V staining and flow cytometry. Cells were stained with FITC Annexin V (BD Biosciences, 556547) and run within 1 hour of staining. 50,000-100,000 events were collected per sample on a CytoFLEX flow cytometer. FlowJo (v10) was used for compensation, gating and analyses of the samples. For cell imaging and cell size analysis, additional replicates were stained in 6-well plates and imaged using a BioTek H1 Synergy instrument (10x objective) and BioTek Gen5 software.

## ACKNOWLEDGEMENTS

The authors thank H. Priode for building the constructs for the apoptosis experiment. This study was supported by grants from the Genentech Research Awards Program (GRAP) and the NIH NCI (R21CA232244) to K. Haynes. This study was supported in part by the following Emory Integrated Core Facilities (EICF): Emory plus Pediatric’s/ Winship Flow Cytometry Core (ECFCC) subsidized by the Emory University School of Medicine, and the Emory Integrated Genomics Core (EIGC) shared resource of Winship Cancer Institute of Emory University and NIH/NCI under award number P30CA138292.

## Notes

https://benchling.com/hayneslab/f_/rmSYkAAU-synthetic-chromatin-actuators-2-0/

### Competing Interest Statement

The authors have declared no competing interest.

## REFERENCES

1. Bernstein, E. et al. Mouse polycomb proteins bind differentially to methylated histone H3 and RNA and are enriched in facultative heterochromatin. Mol. Cell. Biol. 26, 2560–2569 (2006).

2. Lehmann, L. et al. Polycomb repressive complex 1 (PRC1) disassembles RNA polymerase II preinitiation complexes. J. Biol. Chem. 287, 35784–35794 (2012).

3. Kleer, C. G. et al. EZH2 is a marker of aggressive breast cancer and promotes neoplastic transformation of breast epithelial cells. Proc. Natl. Acad. Sci. U. S. A. 100, 11606–11611 (2003).

4. Raaphorst, F. M. et al. Poorly differentiated breast carcinoma is associated with increased expression of the human polycomb group EZH2 gene. Neoplasia 5, 481–488 (2003).

5. Bachmann, I. M. et al. EZH2 expression is associated with high proliferation rate and aggressive tumor subgroups in cutaneous melanoma and cancers of the endometrium, prostate, and breast. J. Clin. Oncol. 24, 268–273 (2006).

6. Chien, Y.-C., Liu, L.-C., Ye, H.-Y., Wu, J.-Y. & Yu, Y.-L. EZH2 promotes migration and invasion of triple-negative breast cancer cells via regulating TIMP2-MMP-2/-9 pathway. Am. J. Cancer Res. 8, 422–434 (2018).

7. Collett, K. et al. Expression of enhancer of zeste homologue 2 is significantly associated with increased tumor cell proliferation and is a marker of aggressive breast cancer. Clin. Cancer Res. 12, 1168–1174 (2006).

8. Holm, K. et al. Global H3K27 trimethylation and EZH2 abundance in breast tumor subtypes. Mol. Oncol. 6, 494–506 (2012).

9. Al-Mahmood, S., Sapiezynski, J., Garbuzenko, O. B. & Minko, T. Metastatic and triple-negative breast cancer: challenges and treatment options. Drug Deliv. Transl. Res. 8, 1483–1507 (2018).

10. Liedtke, C. et al. Response to neoadjuvant therapy and long-term survival in patients with triple-negative breast cancer. J. Clin. Oncol. 26, 1275–1281 (2008).

11. Howard, F. M. & Olopade, O. I. Epidemiology of Triple-Negative Breast Cancer: A Review. Cancer J. 27, 8–16 (2021).

12. Dietze, E. C., Sistrunk, C., Miranda-Carboni, G., O’Regan, R. & Seewaldt, V. L. Triple-negative breast cancer in African-American women: disparities versus biology. Nat. Rev. Cancer 15, 248–254 (2015).

13. Siddharth, S. & Sharma, D. Racial Disparity and Triple-Negative Breast Cancer in African-American Women: A Multifaceted Affair between Obesity, Biology, and Socioeconomic Determinants. Cancers 10, (2018).

14. Lund, M. J. et al. Age/race differences in HER2 testing and in incidence rates for breast cancer triple subtypes: a population-based study and first report. Cancer 116, 2549–2559 (2010).

15. Jene-Sanz, A. et al. Expression of polycomb targets predicts breast cancer prognosis. Mol. Cell. Biol. 33, 3951–3961 (2013).

16. Yomtoubian, S. et al. Inhibition of EZH2 Catalytic Activity Selectively Targets a Metastatic Subpopulation in Triple-Negative Breast Cancer. Cell Rep. 30, 755–770.e6 (2020).

17. King, H. W., Fursova, N. A., Blackledge, N. P. & Klose, R. J. Polycomb repressive complex 1 shapes the nucleosome landscape but not accessibility at target genes. Genome Res. 28, 1494–1507 (2018).

18. Hodges, H. C. et al. Dominant-negative SMARCA4 mutants alter the accessibility landscape of tissue-unrestricted enhancers. Nat. Struct. Mol. Biol. 25, 61–72 (2018).

19. Lehmann, B. D. et al. Multi-omics analysis identifies therapeutic vulnerabilities in triple-negative breast cancer subtypes. Nat. Commun. 12, 6276 (2021).

20. Lee, M.-S., Lim, K., Lee, M.-K. & Chi, S.-W. Structural Basis for the Interaction between p53 Transactivation Domain and the Mediator Subunit MED25. Molecules 23, (2018).

21. Cancer Genome Atlas Network. Comprehensive molecular portraits of human breast tumours. Nature 490, 61–70 (2012).

22. Wasielewski, M., Elstrodt, F., Klijn, J. G. M., Berns, E. M. J. J. & Schutte, M. Thirteen new p53 gene mutants identified among 41 human breast cancer cell lines. Breast Cancer Res. Treat. 99, 97–101 (2006).

23. Kia, S. K., Gorski, M. M., Giannakopoulos, S. & Verrijzer, C. P. SWI/SNF mediates polycomb eviction and epigenetic reprogramming of the INK4b-ARF-INK4a locus. Mol. Cell. Biol. 28, 3457–3464 (2008).

24. Wang, L. et al. The BRG1- and hBRM-associated factor BAF57 induces apoptosis by stimulating expression of the cylindromatosis tumor suppressor gene. Mol. Cell. Biol. 25, 7953–7965 (2005).

25. Tekel, S. J. et al. Tandem Histone-Binding Domains Enhance the Activity of a Synthetic Chromatin Effector. ACS Synth. Biol. 7, 842–852 (2018).

26. Vojnic, E. et al. Structure and VP16 binding of the Mediator Med25 activator interaction domain. Nat. Struct. Mol. Biol. 18, 404–409 (2011).

27. Malik, S. & Roeder, R. G. The metazoan Mediator co-activator complex as an integrative hub for transcriptional regulation. Nat. Rev. Genet. 11, 761–772 (2010).

28. Nyer, D. B., Daer, R. M., Vargas, D., Hom, C. & Haynes, K. A. Regulation of cancer epigenomes with a histone-binding synthetic transcription factor. NPJ Genom Med 2, (2017).

29. Olney, K. C., Nyer, D. B., Vargas, D. A., Wilson Sayres, M. A. & Haynes, K. A. The synthetic histone-binding regulator protein PcTF activates interferon genes in breast cancer cells. BMC Syst. Biol. 12, 83 (2018).

30. McCabe, M. T. et al. EZH2 inhibition as a therapeutic strategy for lymphoma with EZH2-activating mutations. Nature 492, 108–112 (2012).

31. Verma, S. K. et al. Identification of Potent, Selective, Cell-Active Inhibitors of the Histone Lysine Methyltransferase EZH2. ACS Med. Chem. Lett. 3, 1091–1096 (2012).

32. Buchwald, G. et al. Structure and E3-ligase activity of the Ring-Ring complex of polycomb proteins Bmi1 and Ring1b. EMBO J. 25, 2465–2474 (2006).

33. Li, Z. et al. Structure of a Bmi-1-Ring1B polycomb group ubiquitin ligase complex. J. Biol. Chem. 281, 20643–20649 (2006).

34. Gray, F. et al. BMI1 regulates PRC1 architecture and activity through homo- and hetero-oligomerization. Nat. Commun. 7, 13343 (2016).

35. Kreso, A. et al. Self-renewal as a therapeutic target in human colorectal cancer. Nat. Med. 20, 29–36 (2014).

36. Nishida, Y. et al. The novel BMI-1 inhibitor PTC596 downregulates MCL-1 and induces p53-independent mitochondrial apoptosis in acute myeloid leukemia progenitor cells. Blood Cancer J. 7, e527 (2017).

37. Stuckey, J. I. et al. A cellular chemical probe targeting the chromodomains of Polycomb repressive complex 1. Nat. Chem. Biol. 12, 180–187 (2016).

38. Lehmann, B. D. et al. Identification of human triple-negative breast cancer subtypes and preclinical models for selection of targeted therapies. J. Clin. Invest. 121, 2750–2767 (2011).

39. Zhao, M., Kim, P., Mitra, R., Zhao, J. & Zhao, Z. TSGene 2.0: an updated literature-based knowledgebase for tumor suppressor genes. Nucleic Acids Res. 44, D1023–31 (2016).

40. Mahara, S. et al. HIFI-α activation underlies a functional switch in the paradoxical role of Ezh2/PRC2 in breast cancer. Proc. Natl. Acad. Sci. U. S. A. 113, E3735–44 (2016).

41. Shields, C. E., Schnepp, R. W. & Haynes, K. A. Differential Epigenetic Effects of BMI Inhibitor PTC-028 on Fusion-Positive Rhabdomyosarcoma Cell Lines from Distinct Metastatic Sites. Regenerative Engineering and Translational Medicine (2022) doi:10.1007/s40883-021-00244-9.

42. Shields, C. E. et al. Epigenetic regulator BMI1 promotes alveolar rhabdomyosarcoma proliferation and constitutes a novel therapeutic target. Mol. Oncol. 15, 2156–2171 (2021).

43. Ernst, J. et al. Mapping and analysis of chromatin state dynamics in nine human cell types. Nature 473, 43–49 (2011).

44. Thapa, R. J. et al. Interferon-induced RIP1/RIP3-mediated necrosis requires PKR and is licensed by FADD and caspases. Proc. Natl. Acad. Sci. U. S. A. 110, E3109–18 (2013).

45. Kaiser, W. J. et al. Toll-like receptor 3-mediated necrosis via TRIF, RIP3, and MLKL. J. Biol. Chem. 288, 31268–31279 (2013).

46. Seifert, L. & Miller, G. Molecular Pathways: The Necrosome-A Target for Cancer Therapy. Clin. Cancer Res. 23, 1132–1136 (2017).

47. Han, J., Goldstein, L. A., Hou, W. & Rabinowich, H. Functional linkage between NOXA and Bim in mitochondrial apoptotic events. J. Biol. Chem. 282, 16223–16231 (2007).

48. LaBaer, J. et al. New functional activities for the p21 family of CDK inhibitors. Genes Dev. 11, 847–862 (1997).

49. Ding, W. et al. Epigenetic silencing of ITGA2 by MiR-373 promotes cell migration in breast cancer. PLoS One 10, e0135128 (2015).

50. Geoffroy, M., Kleinclauss, A., Kuntz, S. & Grillier-Vuissoz, I. Claudin 1 inhibits cell migration and increases intercellular adhesion in triple-negative breast cancer cell line. Mol. Biol. Rep. 47, 7643–7653 (2020).

51. Adorno-Cruz, V. et al. ITGA2 promotes expression of ACLY and CCND1 in enhancing breast cancer stemness and metastasis. Genes Dis 8, 493–508 (2021).

52. Kowarz, E., Löscher, D. & Marschalek, R. Optimized Sleeping Beauty transposons rapidly generate stable transgenic cell lines. Biotechnol. J. 10, 647–653 (2015).

53. Nath, A. et al. Palmitate-Induced IRE1-XBP1-ZEB Signaling Represses Desmoplakin Expression and Promotes Cancer Cell Migration. Mol. Cancer Res. 19, 240–248 (2021).

54. Zhao, D. et al. TFPI2 suppresses breast cancer progression through inhibiting TWIST-integrin α5 pathway. Mol. Med. 26, 27 (2020).

55. Stephen, H. M. et al. Epigenetic suppression of neprilysin regulates breast cancer invasion. Oncogenesis 5, e207 (2016).

56. Shimo, T. et al. Pathogenic role of connective tissue growth factor (CTGF/CCN2) in osteolytic metastasis of breast cancer. J. Bone Miner. Res. 21, 1045–1059 (2006).

57. Kaustov, L. et al. Recognition and specificity determinants of the human cbx chromodomains. J. Biol. Chem. 286, 521–529 (2011).

58. Morin, R. D., Arthur, S. E. & Assouline, S. Treating lymphoma is now a bit EZ-er. Blood Adv 5, 2256–2263 (2021).

59. Hirukawa, A. et al. Targeting EZH2 reactivates a breast cancer subtype-specific anti-metastatic transcriptional program. Nat. Commun. 9, 2547 (2018).

60. Shi, B. et al. Integration of estrogen and Wnt signaling circuits by the polycomb group protein EZH2 in breast cancer cells. Mol. Cell. Biol. 27, 5105–5119 (2007).

61. Lee, S. T. et al. Context-specific regulation of NF-κB target gene expression by EZH2 in breast cancers. Mol. Cell 43, 798–810 (2011).

62. Chan, H. L. et al. Polycomb complexes associate with enhancers and promote oncogenic transcriptional programs in cancer through multiple mechanisms. Nat. Commun. 9, 3377 (2018).

63. Langmead, B. & Salzberg, S. L. Fast gapped-read alignment with Bowtie 2. Nat. Methods 9, 357–359 (2012).

64. Picard. http://broadinstitute.github.io/picard/.

65. Buenrostro, J. D., Giresi, P. G., Zaba, L. C., Chang, H. Y. & Greenleaf, W. J. Transposition of native chromatin for fast and sensitive epigenomic profiling of open chromatin, DNA-binding proteins and nucleosome position. Nat. Methods 10, 1213–1218 (2013).

66. Jung, Y. H. et al. Maintenance of CTCF- and Transcription Factor-Mediated Interactions from the Gametes to the Early Mouse Embryo. Mol. Cell 75, 154–171.e5 (2019).

67. Lyu, X., Rowley, M. J. & Corces, V. G. Architectural Proteins and Pluripotency Factors Cooperate to Orchestrate the Transcriptional Response of hESCs to Temperature Stress. Mol. Cell 71, 940–955.e7 (2018).

68. Jung, Y. H. et al. Chromatin States in Mouse Sperm Correlate with Embryonic and Adult Regulatory Landscapes. Cell Reports vol. 18 1366–1382 Preprint at https://doi.org/10.1016/j.celrep.2017.01.034 (2017).

69. Liu, T. Use Model-Based Analysis of ChIP-Seq (MACS) to Analyze Short Reads Generated by Sequencing Protein–DNA Interactions in Embryonic Stem Cells. in Stem Cell Transcriptional Networks: Methods and Protocols (ed. Kidder, B. L.) 81–95 (Springer New York, 2014).

70. Amemiya, H. M., Kundaje, A. & Boyle, A. P. The ENCODE Blacklist: Identification of Problematic Regions of the Genome. Sci. Rep. 9, 9354 (2019).

71. Robinson, J. T. et al. Integrative genomics viewer. Nat. Biotechnol. 29, 24–26 (2011).

72. Haynes, K. A. & Priode, J. H. Rapid Single-Pot Assembly of Modular Chromatin Proteins for Epigenetic Engineering. Methods Mol. Biol. 2599, 191–214 (2023).

