## Supplemental Data for "PIC recruitment by synthetic reader-actuators to polycomb-silenced genes blocks triple-negative breast cancer invasion"

Figure S1. Polycomb expression is elevated and tumor suppressor genes are downregulated in TNBC patient samples

Figure S2. BT-549 IC50 values for polycomb inhibitors used in this study

Figure S3. Comparison of SRA-UpDEG baseline expression in normal versus TNBC cell lines

Table S1. Nanostring expression data from Figure 2E

Table S2. RNA-seq data for SRA UpDEGs and the 100 top highly expressed genes from Figure 3C

Table S3. SRA UpDEGs and associated KEGG pathways

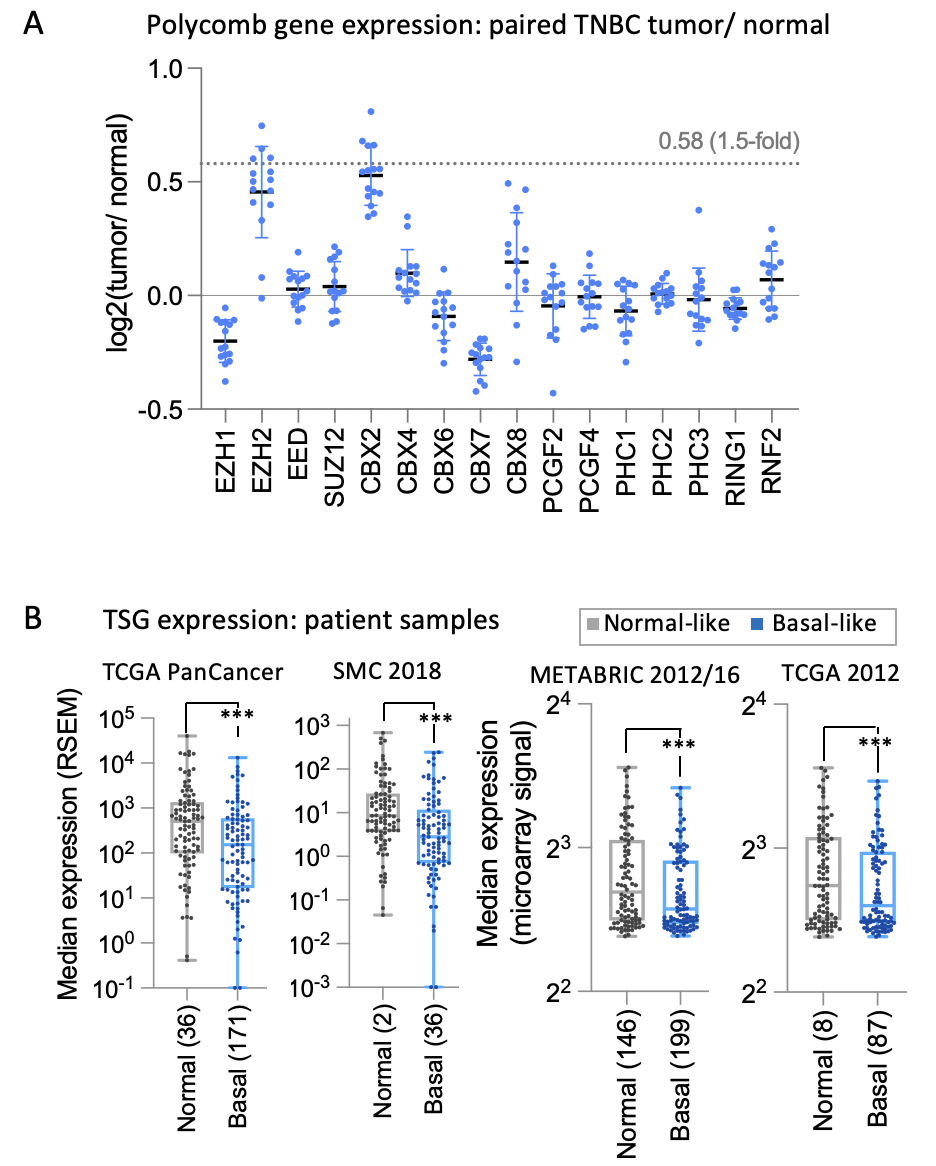

**Figure S1.** Polycomb expression is elevated and tumor suppressor genes are downregulated in TNBC patient samples. (A) Expression levels for polycomb subunit-encoding genes (publicly available RNA-seq data from cBioPortal) is shown for basal-like triple negative tumor versus normal tissue from the same patient (Xena database). (B) Box plots show baseline expression levels (RNA-seq data, cBioPortal) of 97 tumor suppressor genes (out of 589 BRCA TSGs from the TSGene database[1]) that are consistently > 2-fold downregulated in patient samples classified as TNBC/ Basal versus normal. Each point is the median mRNA signal for *n* samples for one gene. In the TCGA PanCancer 2012 study[2] basal samples were 80% TNBC, 10% ER+/HER2−, and 2% HER2+. TNBC types were a small fraction of luminal A, luminal B, and Her2 groups (2%, 1%, and 9% respectively). Two-tailed paired t-test *p* < 0.001***.

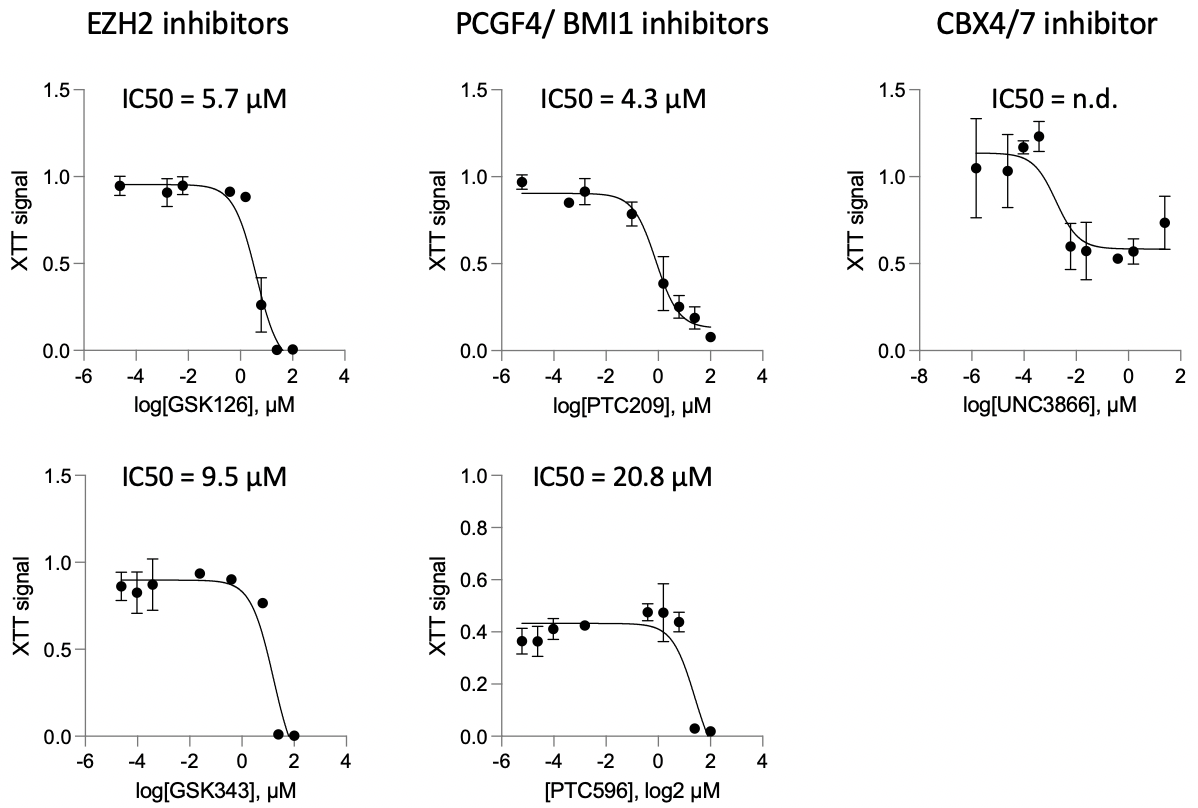

**Figure S2.** BT-549 IC50 values for polycomb inhibitors. Cells were seeded at 1.0E4 per well in a 96-well plate coated with 2% ECM-supplemented medium (Millipore Sigma, E0282). Cells were treated with DMSO or a final concentration of 1.49E-6 to 1.0E2 μM of each inhibitor in 200 µL complete media per well. After 3 days of treatment, XTT Reagent and Activation Reagent (ATCC, 30-1011K) were mixed according to the manufacturer’s instructions, and 50 µL of activated-XTT solution was added to each well. Plate readings were taken at wavelengths of 475 nm (specific) and 660 nm (non-specific background) using a Biotek Cytation 1 plate reader. Plotting of log transformed means with standard errors for triplicate samples, and nonlinear fit to determine IC50 values were done using GraphPad Prism 9 software. No IC50 value for UNC3866 could be determined because viability was reduced by less than 50% at the highest concentration. Therefore, we used a concentration of 30 μM, which has been used in other *in vitro* carcinoma studies[3].

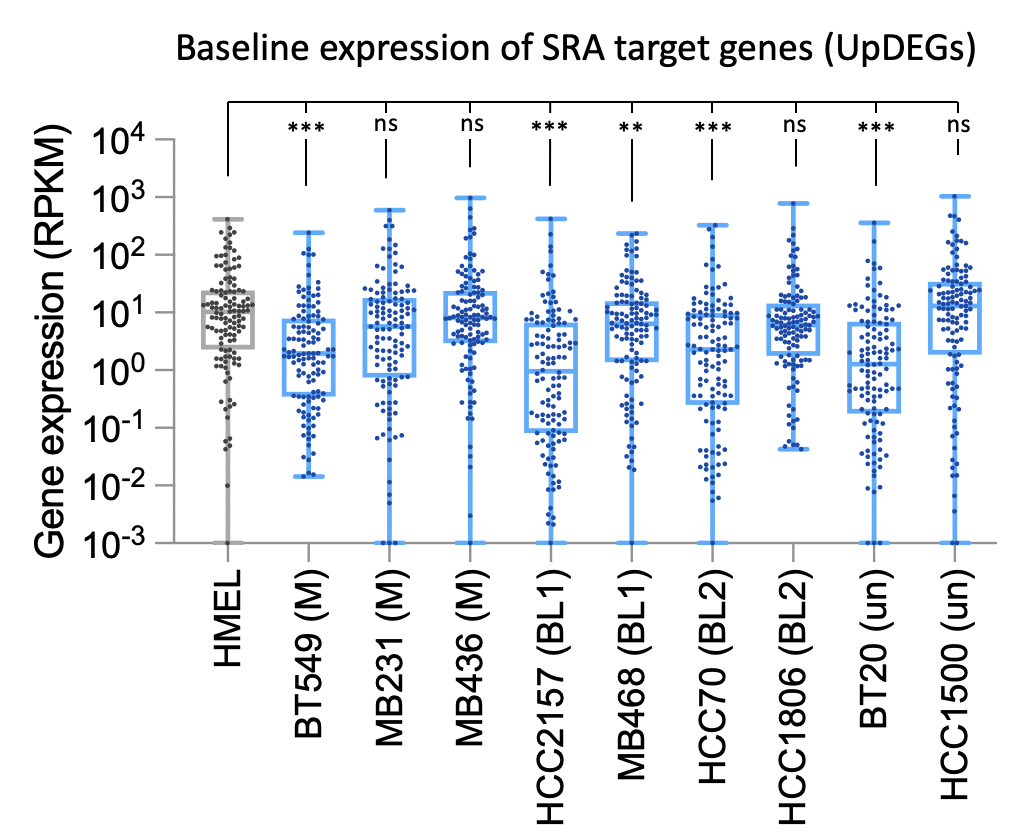

**Figure S3.** Comparison of SRA-UpDEG baseline expression in normal versus TNBC cell lines. Box plots show baseline expression levels (RNA-seq data, cBioPortal) of the 122 protein coding SRA UpDEGs from tumor suppressor genes in patient samples classified as normal or TNBC/ Basal cancer types. Two-tailed paired t-test *p* < 0.001***, *p* < 0.01**, *p* <0.05**.

**Table S1.** Nanostring expression data from Figure 2E. See Excel spreadsheet.

**Table S2.** RNA-seq data for SRA UpDEGs and the 100 top highly expressed genes from Figure 3C. See Excel spreadsheet.

**Table S3.** SRA UpDEGs and associated KEGG pathways from Figure 4

| **Category** | **KEGG Pathway** | **Gene Symbols** |  |
| --- | --- | --- | --- |
|  |  | Agonists | Antagonists |
| Cell death | Necroptosis (hsa04217) | TLR3, STAT1, MLKL, STAT2, IL1A, IL1B | BIRC3, TNFAIP3 |
|  | TNF signaling pathway (hsa04668) | MLKL, IL1B, CXCL10 | BIRC3, TNFAIP3 |
|  | Apoptosis (hsa04210) | PMAIP1 | BIRC3 |
| Cell cycle arrest | Cellular senescence (hsa04218) | CDKN1A, HLA-B, HLA-E, HLA-C, HLA-A, IL1A |  |
| Migration & invasion block | ECM-receptor interaction (hsa04512) | ITGA2 |  |
|  | Tight junction (hsa04530) | CLDN1 |  |
| Immune surveillance | Herpes simplex virus 1 infection (hsa05168) | TLR3, STAT1, IFNB1, SP100, IFIH1, TAP1, HLA-B, HLA-E, STAT2, HLA-C, HLA-A, B2M, OAS1, BST2, IL1B, MYD88, OAS2, CCL5, OAS3, C3 | BIRC3 |
|  | NOD-like receptor signaling pathway (hsa04621) | GBP4, STAT1, IFNB1, GBP1, GBP3, STAT2, OAS1, GBP5, TNFAIP3, IL1B, MYD88, BIRC3, OAS2, CCL5, OAS3 |  |
|  | Toll-like receptor signaling pathway (hsa04620) | TLR3, STAT1, IFNB1, IL1B, CXCL10, MYD88 |  |
|  | Antigen processing and presentation (hsa04612) | TAP1, HLA-B, HLA-E, HLA-C, HLA-A, B2M |  |
|  | JAK-STAT signaling pathway (hsa04630) | STAT1, PI3, STAT2, CSF3 |  |
|  | Cytokine-cytokine receptor interaction (hsa04060) | GDF15, IL1B, CXCL10 |  |
|  | Complement and coagulation cascades (hsa04610) | SERPING1, SERPINE2 |  |
|  | B cell receptor signaling pathway (hsa04662) | IFITM1 |  |
| Cancer progression | Breast cancer (hsa05224) | DLL1, LEF1 | CDKN1A |
|  | Wnt signaling pathway (hsa04310) | LEF1 |  |
|  | B cell receptor signaling pathway (hsa04662) | IFITM1 |  |

**Supplemental References**

1. Zhao, M., Kim, P., Mitra, R., Zhao, J. & Zhao, Z. TSGene 2.0: an updated literature-based knowledgebase for tumor suppressor genes. *Nucleic Acids Res*. **44**, D1023–31 (2016).
2. Cancer Genome Atlas Network. Comprehensive molecular portraits of human breast tumours. *Nature* **490**, 61–70 (2012).
3. Stuckey, J. I. et al. A cellular chemical probe targeting the chromodomains of Polycomb repressive complex 1. *Nat. Chem. Biol.* **12**, 180–187 (2016).
